# Increased atherosclerosis and expression of inflammarafts in macrophage foam cells in AIBP-deficient mice

**DOI:** 10.1101/2025.08.12.669996

**Authors:** Shenglin Li, Nicolaus Nazarenkov, Elena Alekseeva, Juliana Maria Navia-Pelaez, Aakash Patel, Patrick Secrest, Philip L.S.M. Gordts, Sven Heinz, Yury I. Miller

## Abstract

Atherosclerotic lesions comprise different populations of macrophages, including lipid-laden macrophage foam cells and non-foamy, inflammatory macrophages, which play distinct roles in disease progression. Non-foamy macrophages express higher levels of inflammarafts – enlarged, cholesterol-rich lipid rafts hosting assemblies of inflammatory receptors – compared to foam cells in atherosclerotic lesions of *Ldlr^−/−^*mice. Apolipoprotein A-I binding protein (AIBP) has been shown to control lipid raft dynamics. This study investigated the effect of systemic AIBP deficiency on inflammaraft expression in foam cells and non-foamy macrophages in atherosclerotic lesions of hypercholesterolemic mice. A larger number of foam cells, with increased neutral lipid accumulation, populated atherosclerotic lesions in *Apoa1bp^−/−^Ldlr^−/−^*mice compared to *Ldlr^−/−^* mice. Importantly, AIBP-deficient foam cells expressed higher levels of TLR4 dimers and lipid rafts (markers of inflammarafts) than control mice, accompanied by larger atherosclerotic lesions and larger necrotic cores compared to *Ldlr^−/−^*mice. In a model of foam cells, *Apoa1bp^−/−^* bone marrow-derived macrophages incubated with oxidized LDL had increased expression of inflammation- and atherosclerosis-related genes. These results indicate that AIBP deficiency is associated with a proinflammatory transition of foam cells in which increased lipid accumulation is paradoxically associated with an increased expression of inflammarafts and correlates with the development of advanced atherosclerotic plaques.

## Introduction

The advances in flow cytometry and sequencing techniques have helped identify distinct populations of macrophages in human and mouse atherosclerotic lesions.^[1,2]^ One differentiating feature in lesional macrophages is the handling of excess cholesterol. Lipid-laden macrophage foam cells accumulate large deposits of esterified cholesterol in lipid droplets. In contrast, non-foamy macrophages display increased levels of unesterified cholesterol-rich lipid rafts in the plasma membrane, which, due to low diffusion rates and the presence of cholesterol and sphingolipid binding domains in many proteins, serve as a platform for the assembly of functional receptor complexes.^[3]^ Enlarged lipid rafts that host assembled complexes of inflammatory receptors have been designated as inflammarafts.^[4]^ In atherosclerotic lesions, non-foamy macrophages, which express inflammatory genes^[1]^, retain high levels of inflammarafts, particularly inflammarafts rich in TLR (Toll-like receptor) −4 dimers.^[3]^ In contrast, macrophage foam cells express lipid storage genes, have low expression of inflammatory genes^[1]^ and low abundance of TLR4 inflammarafts.^[3]^ However, a recent single-cell RNA-seq study has identified in human carotid lesions a population of macrophages expressing high levels of *PLIN2*, a lipid droplet protein, and TLR-driven inflammatory genes.^[5]^ The mechanisms regulating the emergence of inflammatory lipid-laden macrophages remain to be fully elucidated.

One possible mechanism may involve the function of apolipoprotein A-I binding protein (AIBP, gene name *Apoa1bp* or *Naxe*), which facilitates cholesterol efflux by binding to apoA-I and stabilization of ATP-binding cassette transporter A1 (ABCA1). The removal of cholesterol from the plasma membrane reduces the abundance of TLR4 inflammarafts.^[6–8]^ AIBP is ubiquitously expressed and can be secreted in response to apoA-I or oxidized LDL (OxLDL).^[9,10]^ Human and mouse atherosclerotic lesions express high levels of AIBP protein. ^[10,11]^ *Apoa1bp^−/−^* mice fed a Western-type diet exhibit rapid weight gain and impaired glucose tolerance, hallmarks of metabolic disease.^[12]^ Intracellular AIBP localizes to mitochondria and regulates mitophagy.^[10,11]^ *Apoa1bp^−/−^Ldlr^−/−^* mice fed a high-cholesterol, high-fat diet have more atherosclerosis compared to *Ldlr^−/−^* mice. ^[12]^ In mouse models of neuropathic pain and Alzheimer’s disease, *Apoa1bp^−/−^* mice have chronic, increased expression of TLR4 inflammarafts in spinal cord or brain microglia, respectively, accompanied by oxidative stress and mitochondrial dysfunction. ^[13,14]^

Here, we demonstrate that systemic AIBP deficiency in hyperlipidemic mice is associated with a proinflammatory transition of macrophage foam cells, characterized by increased lipid accumulation and increased expression of TLR4 inflammarafts, as well as larger necrotic cores in atherosclerotic lesions.

## Results

### Increased inflammarafts in lesional macrophage foam cells in AIBP-deficient mice

In cross-sections of the aortic root of *Apoa1bp^−/−^Ldlr^−/−^* and *Ldlr^−/−^* mice fed a 16-week high-fat diet (Figure 1A), AIBP-deficient male mice showed a higher percentage of F4/80^+^ area within the LipidTOX^+^ neutral lipid positive lesion area, than *Ldlr^−/−^*male mice (Figure 1B and 1C), indicating that more macrophages populated the atherosclerotic lesions in AIBP-deficient mice. Next, aortic single-cell suspensions were gated for BODIPY-high (foamy) and BODIPY-low (non-foamy), CD45^+^F4/80^+^ macrophages (Supplemental Figures S1 and S2). The proportion of macrophage foam cells was significantly increased in *Apoa1bp^−/−^Ldlr^−/−^* mice (foamy/non-foamy = 7:3) when compared to *Ldlr^−/−^* mice (4:6) (Figure 1D and 1F). The macrophage foam cells from *Apoa1bp^−/−^Ldlr^−/−^* mice had more neutral lipid accumulated, measured by the BODIPY intensity, than *Ldlr^−/−^*foam cells (Figure 1E and 1G).

**Figure 1.**
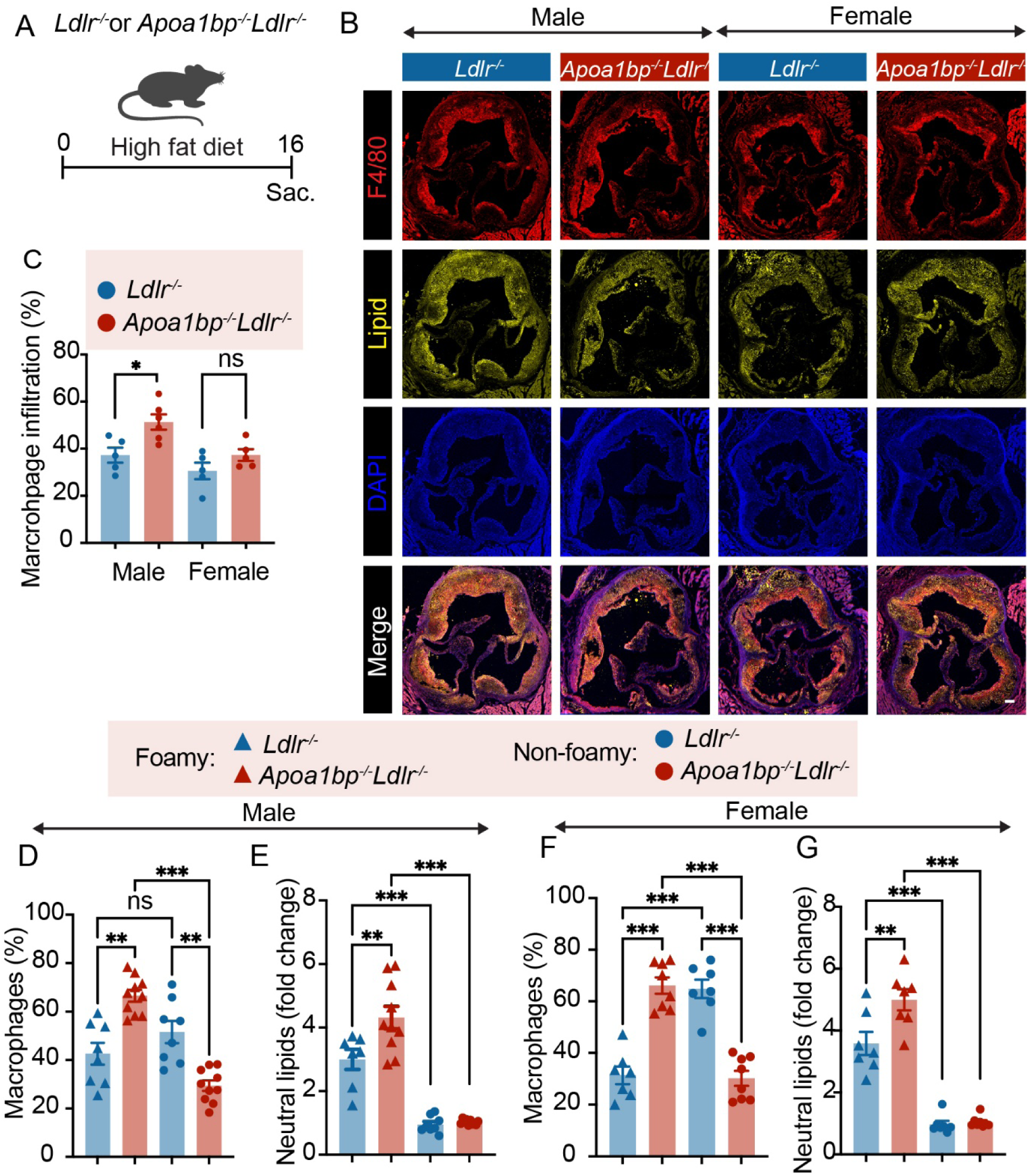
Higher numbers of foam cells with higher levels of neutral lipids in AIBP-deficient mice. **A.** Experimental design of a high-fat diet feeding experiment. **B.** Representative images of immunostaining of aortic root. Scale bar = 100 μm. **C.** Quantitative analysis of F4/80^+^ area in **B**. **D** and **E.** Data from male mice (n=8-10). **F** and **G.** Data from female mice (n=7-8). **D/F.** Percentages of foam cells and non-foamy macrophages in atherosclerotic lesions from *Apoa1bp^−/−^Ldlr^−/−^*and *Ldlr^−/−^* mice. **E/G.** Content of neutral lipids in foam cells, normalized to the readings in non-foamy macrophages from the same aorta. One-way ANOVA with Tukey’s. Mean ± SEM; ns, nonsignificant; *, *p* ≤0.05; **, *p* ≤0.01; ***, *p* ≤0.001.

We then measured the degree of TLR4 dimerization in lesional macrophages as a key parameter of TLR4 inflammaraft expression. The expression of TLR4 dimers in aortic macrophages was significantly higher than that in splenic macrophages stimulated with LPS, used as an intra-assay positive control (Figure 2A). Further, both macrophage foam cells and non-foamy macrophages from *Apoa1bp^−/−^Ldlr^−/−^*mice expressed significantly higher TLR4 dimers, when compared to those from *Ldlr^−/−^* mice (Figure 2A and 2D). The difference was further exacerbated when TLR4 dimers were integrated over the number (percentage) of each macrophage population, showing that macrophage foam cells from *Apoa1bp^−/−^Ldlr^−/−^* male and female mice were the predominant population expressing TLR4 dimers (Figure 2B and 2E). In contrast, in *Ldlr^−/−^* mice, the major TLR4 dimer-expressing cells were non-foamy macrophages (Figure 2B and 2E). Furthermore, *Apoa1bp^−/−^Ldlr^−/−^* macrophage foam cells exhibited increased levels of lipid rafts, equal to the lipid raft levels in non-foamy macrophages in both *Apoa1bp^−/−^Ldlr^−/−^* and *Ldlr^−/−^*mice (Figure 2C and 2F). These results indicate that AIBP deficiency leads to increased TLR4 inflammaraft expression in macrophage foam cells, making them the major inflammaraft-expressing subpopulation in atherosclerotic plaques in AIBP-deficient mice (Figure 2G).

**Figure 2.**
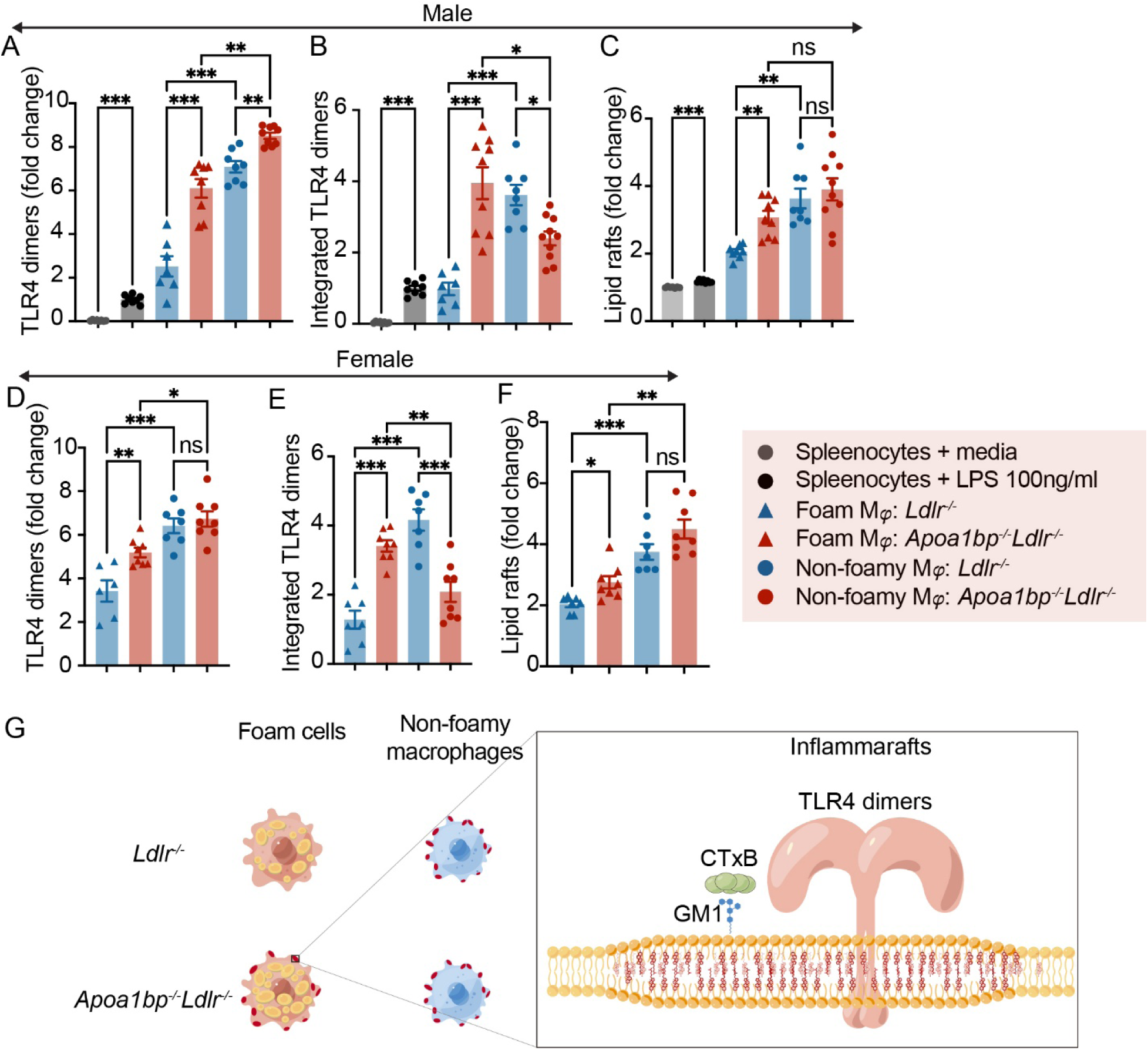
Foam cells are the major macrophage population expressing TLR4 inflammarafts in AIBP-deficient mice. **A-C.** Data from male mice (n=8-10). **D-F.** Data from female mice (n=7-8). **A/D.** Expression of TLR4 dimers in foam cells and non-foamy macrophages in *Apoa1bp^−/−^Ldlr^−/−^* and *Ldlr^−/−^* mice. Splenocytes stimulated with vehicle or LPS (100 ng/ml, 15 min) served as negative and positive controls in **A-F**. **B/E**. Integrated TLR4 dimers (TLR4 dimers in each cell type in **A/D** multiplied by cell type percentage in Figure 1D and 1F. **C/F**. Expression of lipid rafts in lesional macrophages. **G.** Illustration of a foam cell phenotype in *Apoa1bp^−/−^Ldlr^−/−^*mice characterized by higher lipid accumulation and increased expression of TLR4 inflammarafts. GM1: monosialotetrahexosylganglioside. CTxB: cholera toxin subunit B. One-way ANOVA with Tukey’s. Mean ± SEM; ns, nonsignificant; *, *p* ≤0.05; **, *p* ≤0.01; ***, *p* ≤0.001.

To model macrophage foam cells, we incubated bone marrow-derived macrophages (BMDM), obtained from *Apoa1bp^−/−^Ldlr^−/−^* or *Ldlr^−/−^* mice, with 20 μg/mL OxLDL for 24 hours. A total of 604 genes were upregulated, and 660 genes were downregulated in *Apoa1bp^−/−^Ldlr^−/−^*or OxLDL-treated BMDMs compared to those from *Ldlr^−/−^* mice (Figure 3A). Among the top differentially expressed genes were *Wdfy1*, suggesting a predominant TRIF signaling downstream of TLR4^[15]^, *P2ry12*, a major purinergic receptor in macrophages mediating vascular inflammation^[16]^, as well as *Mmp9* and several genes encoding extracellular matrix proteins, suggesting vulnerable lesion remodeling^[17]^ (Figure 3B).

**Figure 3.**
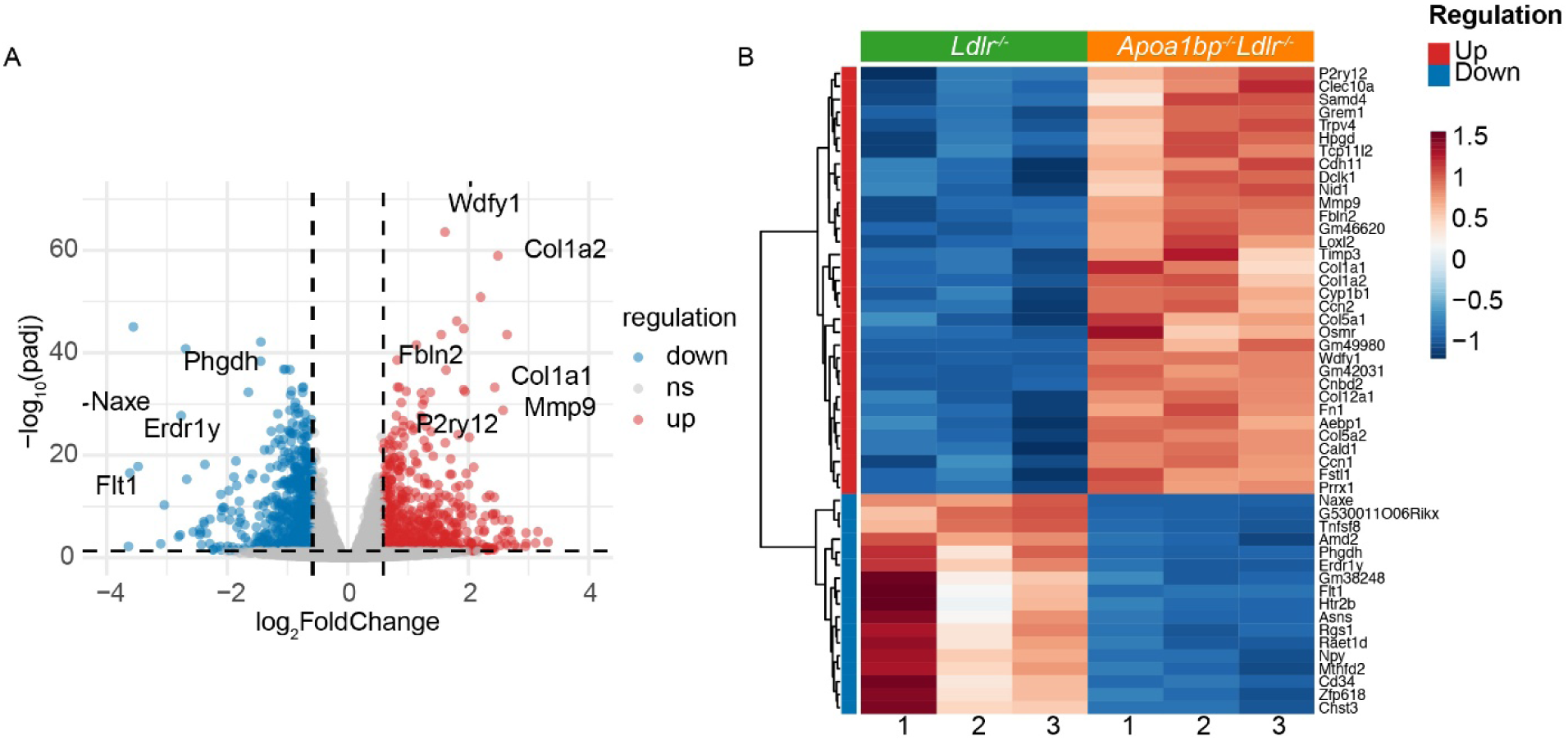
Transcriptional changes in foamy macrophage cell models of *Apoa1bp^−/−^Ldlr^−/−^*and *Ldlr^−/−^*mice. Bone marrow-derived macrophages (BMDM) from *Apoa1bp^−/−^Ldlr^−/−^* and *Ldlr^−/−^*male mice were incubated with 20 μg/mL OxLDL for 24 hours to mimic macrophage foam cells. **A**. Volcano plot **B.** Heatmap showing the top 50 differentially expressed genes selected based on a combined rank of adjusted p-value (ascending) and absolute log2 fold-change (descending).

### Advanced atherosclerosis in AIBP-deficient mice

To assess the extent of atherosclerosis, consecutive cross-sections were collected from the aortic root for quantification. Larger atherosclerotic lesions were observed in the proximal region of the aortic valve in *Apoa1bp^−/−^Ldlr^−/−^* male mice when compared to *Ldlr^−/−^* mice (Figure 4A and 4B). AIBP-deficient female mice displayed a larger distal (500 to 600 µm from valve origin) lesion size (Figure 4A and 4C). The analysis of total lesion volume indicated a tendency for increased atherosclerotic plaque burden (Figure 4D). We observed sex differences in the atherosclerotic lesion morphology in *Apoa1bp^−/−^Ldlr^−/−^* mice (Figure 4B and 4C). In *Ldlr^−/−^* mice, the lesion size curves in both males and females were bell-shaped, peaking at 400 μm from the aortic root. In *Apoa1bp^−/−^Ldlr^−/−^*mice, the lesion size curve in female mice continuously ascended all the way from the valve origin to the 600 μm distance, while the curve in male mice remained bell-shaped, peaking at 500 μm.

**Figure 4.**
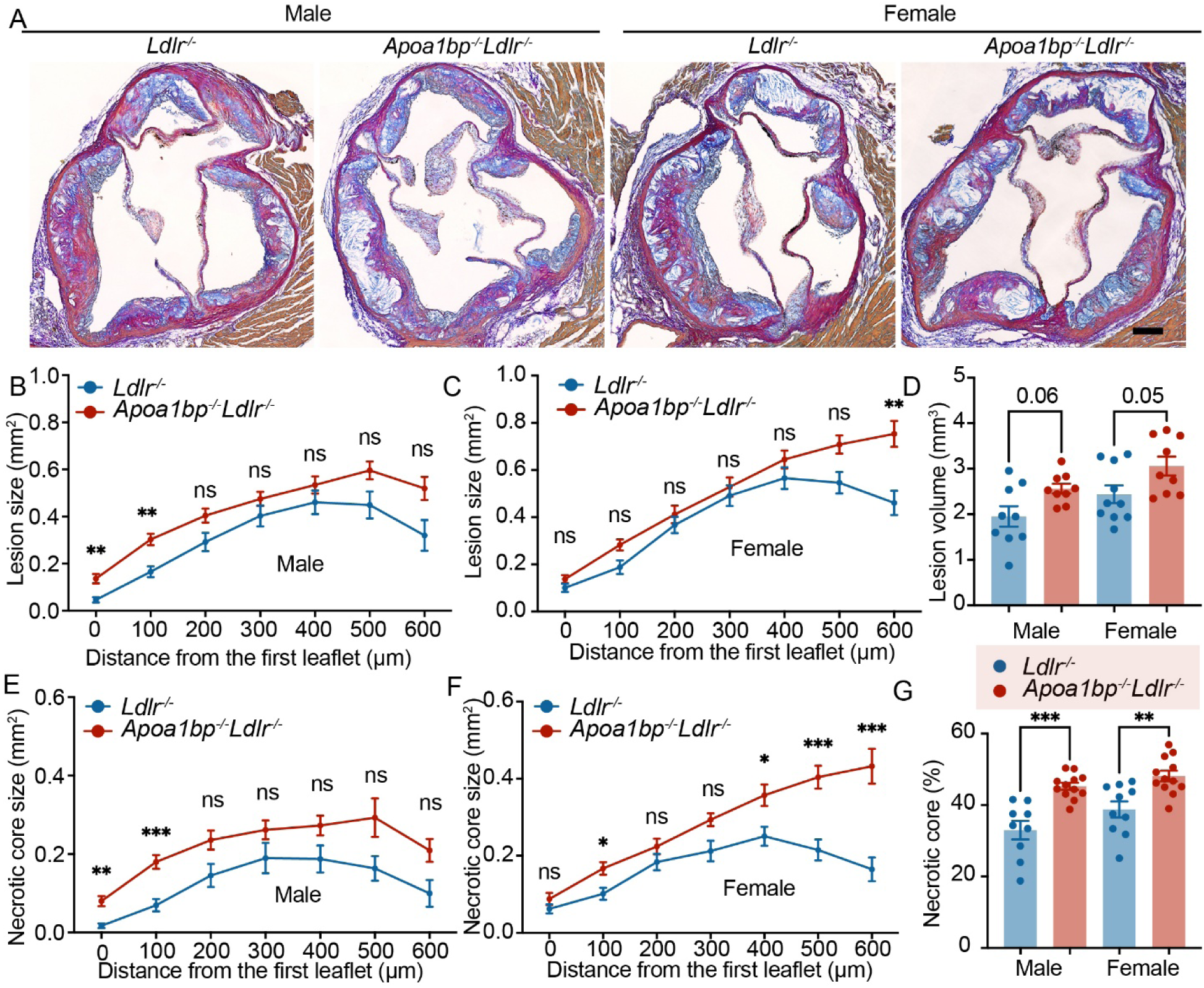
Advanced atherosclerotic lesions in AIBP-deficient mice. **A.** Representative images of the aortic root stained with modified Van Gieson staining. Scale bar = 100 μm. Quantitative analysis of lesion size in male mice (**B**) and female mice (**C**). **D**. Lesion volume measured by area under the curve from **B** and **C**. Quantitative analysis of necrotic core size in male mice (**E**) and female mice (**F**). **G.** Quantitative analysis of necrotic core percentage over the total lesion area. N = 9-10; two-way ANOVA with Tukey’s or Dunnett’s multiple comparison tests (B, C, E, F), and t-test (D and G). Mean ± SEM; ns, nonsignificant; *, *p* ≤0.05; **, *p* ≤≦0.01; ***, *p* ≤0.001.

Formation and enlargement of a necrotic core imply plaque destabilization, associated with vascular inflammation, and indicate advanced atherosclerotic plaques.^[18,19]^ Consistent with the lesion size measurements, a larger necrotic core size was observed in the proximal region of the aortic valve in *Apoa1bp^−/−^Ldlr^−/−^* male mice (Figure 4A and 4E) and at the distal region of the aortic valve in *Apoa1bp^−/−^Ldlr^−/−^*female mice (Figure 4A and 4F) when compared to *Ldlr^−/−^* mice. Of note, the percentage of necrotic core in the lesion area was significantly higher in *Apoa1bp^−/−^Ldlr^−/−^*mice than in *Ldlr^−/−^* mice, in both male and female, suggesting a more vulnerable advanced plaque in AIBP-deficient mice (Figure 4G).

*Apoa1bp^−/−^Ldlr^−/−^* mice gained body weight faster than *Ldlr^−/−^* mice (Figure 5A and 5B) and had higher plasma triglyceride but not cholesterol levels (Figure 5C and 5D). Overall, male and female mice showed similar results. These results indicate that AIBP deficiency affects lipid homeostasis and leads to a larger lesion burden and advanced atherosclerotic plaques.

**Figure 5.**
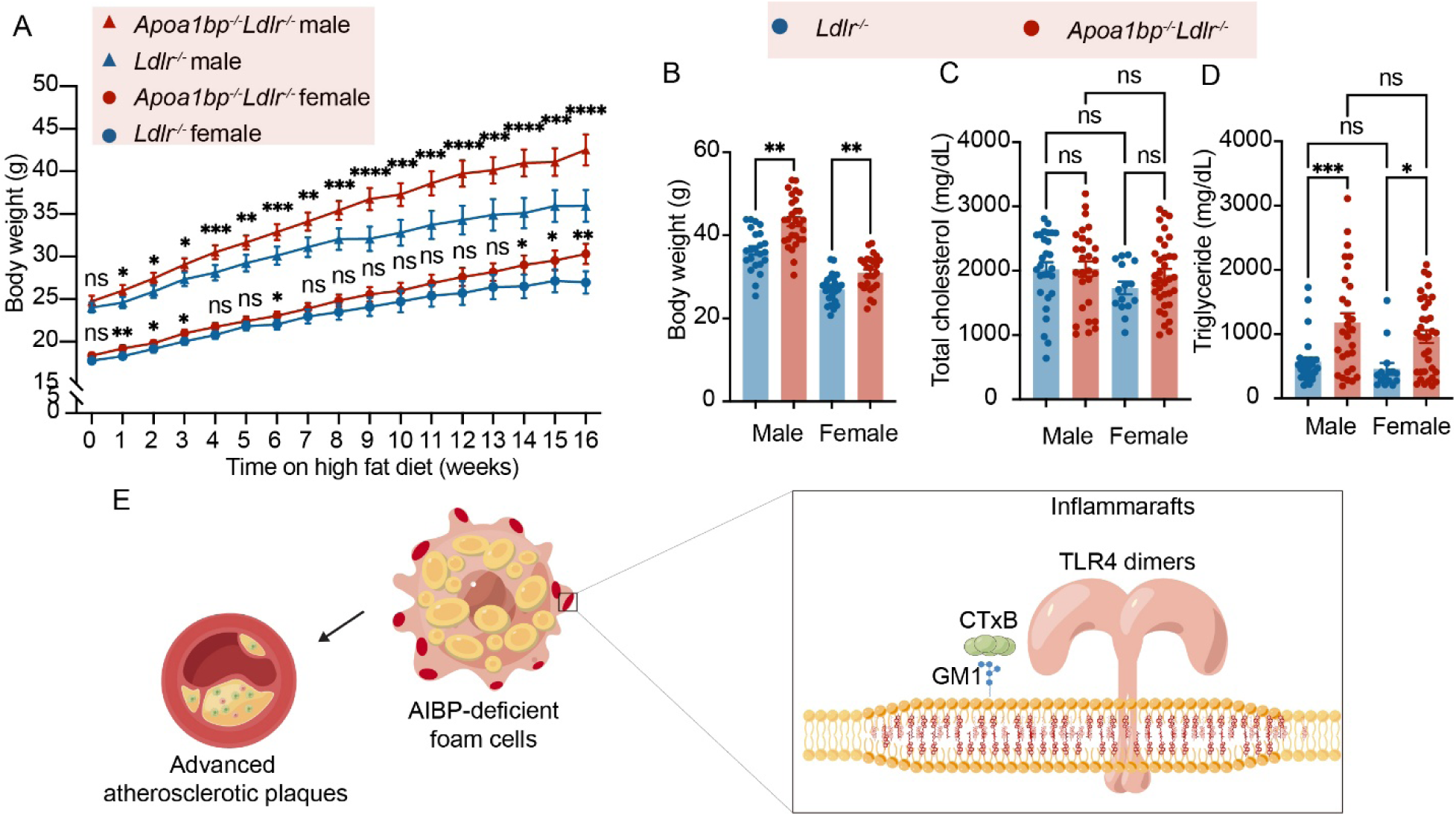
Weight gain and plasma lipid levels. **A.** Body weight gain of *Apoa1bp^−/−^Ldlr^−/−^*and *Ldlr^−/−^* mice during HFD feeding. **B.** End point body weight. **C.** Plasma total cholesterol levels. **D.** Plasma triglyceride levels. **E.** Diagram illustrating this paper’s findings: AIBP deficiency leads to marked expression of inflammarafts and increased lipid accumulation in lesional macrophage foam cells. Advanced atherosclerotic plaques accompany the transition of foam cells to a proinflammatory phenotype. N = 23-29; two-way and one-way ANOVA with Tukey’s or Dunnett’s multiple comparison tests for statistical analysis. Results are presented as Mean ± SEM; ns, nonsignificant; *, *p* ≤0.05; **, *p* ≤0.01; ***, *p* ≤0.001.

## Discussion

In this study, we demonstrated that the atheroprotective effect of AIBP is strongly associated with the attenuation of TLR4 inflammarafts in macrophage foam cells. AIBP deficiency leads to marked expression of inflammarafts and increased lipid accumulation in lesional macrophage foam cells. Advanced atherosclerotic plaques accompanied the transition of foam cells to a proinflammatory phenotype.

Multiple lines of evidence support the concept that macrophage foam cells express lipid storage genes but exhibit suppressed expression of inflammatory-response genes, leading to the reduced contribution of foam cells to inflammation in atherosclerosis compared to non-foamy macrophages. Macrophage foam cells from *Ldlr^−/−^* mice exhibit altered cholesterol metabolism and a marked suppression of proinflammatory mediators that are normally characteristic of the inflammatory responses associated with atherosclerotic lesions.^[20]^ Similarly, transcriptome data from *Apoe^−/−^* mice show that lipid-rich macrophage foam cells express few inflammatory genes but many lipid-processing genes, while non-foamy macrophages are the primary population expressing IL-1β and other inflammatory mediators in the atherosclerotic aorta.^[1]^ Moreover, we previously established that non-foamy macrophages in atherosclerotic lesions of *Ldlr^−/−^* mice express higher levels of inflammarafts than macrophage foam cells, correlating with increased plasma levels of inflammatory cytokines such as monocyte chemoattractant protein-1 (MCP-1), interleukin (IL)-6, and IL-1β. ^[3]^

Despite the evidence for a non-inflammatory character of macrophage foam cells, the mechanisms by which macrophage foam cells in atherosclerotic lesions exhibit less inflammatory response than other macrophage populations remain poorly understood. The most notable difference between foam cells and non-foamy macrophages is the content of intracellular lipid droplets, as well as distinct lipid metabolism. High levels of desmosterol, the last intermediate in the Bloch pathway of cholesterol biosynthesis, have been reported to be associated with suppression of proinflammatory mediators in macrophage foam cells via several homeostatic pathways, including activation of LXR target genes, inhibition of SREBP target genes, and selective reprogramming of fatty acid metabolism.^[20]^ These findings indicate that lipid metabolism is involved in the suppression of inflammatory mediators. On the other hand, a recent publication found the TLR-dependent pathogenic macrophage transition to an inflammatory lipid-associated phenotype in human intraplaque immune cell communities.^[5]^

AIBP not only facilitates cholesterol efflux from the cell membrane to regulate lipid metabolism, but AIBP-mediated disruption of lipid rafts also protects microglia from TLR4 inflammaraft-associated oxidative stress and mitochondrial dysfunction. ^[13]^ In our study, we observed that AIBP deficiency leads to enhanced expression of TLR4 dimers and lipid rafts in macrophage foam cells, as well as increased lipid accumulation in lesional macrophages, resulting in macrophage foam cells being the major inflammaraft-expressing macrophage subpopulation in atherosclerotic lesions of *Apoa1bp^−/−^Ldlr^−/−^* mice. The proinflammatory transition of foam cells accompanied the formation of advanced atherosclerotic plaques. In this study, we examined gene expression in AIBP-deficient and wild-type foam cells using BMDMs incubated with OxLDL. This cellular model only partially replicates the in vivo environment surrounding vascular macrophages in mice with systemic AIBP expression or deficiency, where AIBP is secreted by macrophages to have an autocrine effect^[10]^ and also by other cell types for a paracrine effect. In fact, the choice of a systemic *Apoa1bp^−/−^Ldlr^−/−^* mouse for this study was dictated by the secreted nature of ubiquitously expressed AIBP, which exerts its major effect on lipid rafts extracellularly. Among the upregulated genes in AIBP-deficient BMDMs were the genes broadly associated with atherosclerosis progression, including the pathways not directly downstream of TLR4 signaling. These findings suggest the assembly of other receptor complexes in inflammarafts, as we reported earlier.^[3]^ So far, flow cytometry-based detection of TLR4 dimers and lipid rafts is the most robust measure of inflammarafts, which nevertheless host other, diverse receptor assemblies. Development of more comprehensive methods to characterize inflammarafts in macrophages and other vascular cells will advance the understanding of the inflammatory milieu in atherosclerosis.

Interestingly, our analysis of aortic root lesions showed a sex difference in lesion distribution in response to AIBP deficiency. Female *Apoa1bp^−/−^Ldlr^−/−^*mice exhibited a continuous increase in lesion size and necrotic core size starting from the origin of the aortic valve, while lesions in male *Apoa1bp^−/−^Ldlr^−/−^*mice and both sexes of *Ldlr^−/−^* mice reached the maximum size at 400-500 µm from the origin. These findings indicate that AIBP contributes to the sex difference observed in atherosclerosis. ^[21,22]^

In our previous study with *Apoa1bp^−/−^Ldlr^−/−^*mice, we observed elevated plasma cholesterol and triglyceride levels,^[12]^ in contrast to the results of this work, which showed increased triglyceride but not cholesterol plasma levels. The likely reason for this difference is the difference in the high-fat diets used in these two studies. The Western diet used in the previous study contained 1.25% cholesterol^[12]^, which is higher than the 0.2% cholesterol diet in this study. One limitation of the present study is that the analyses were conducted at one time point, 16 weeks after the start of diet intervention. Without earlier time points, it is unclear whether the proinflammatory macrophage foam cell phenotype represents a stable identity or a transitional stage from non-foamy to foamy macrophages.

The major finding of this study – that AIBP suppresses TLR4 inflammaraft expression specifically in macrophage foam cells – warrants a detailed, genome-wide analysis of gene expression in foamy and non-foamy macrophages from atherosclerotic lesions of *Apoa1bp^−/−^Ldlr^−/−^* and *Ldlr^−/−^*mice. This will help elucidate the mechanisms by which AIBP regulates inflammarafts and the signaling pathways in macrophage foam cells that prevent them from assuming an inflammatory phenotype.

## Methods

### Animals

*Apoa1bp^−/−^Ldlr^−/−^* mice^[12]^ and *Ldlr^−/−^* mice were bred in-house and housed at a maximum of five animals of the same sex per cage at room temperature and a 12:12 light-dark cycle. Age- and weight-matched mice were used. Both male and female mice were included. Starting at eight weeks of age, mice were fed a 16-week Western-style diet containing 42% kcal from fat and 0.2% cholesterol (TD.88137, Envigo Teklad). Mice were fasted for 8 hours before euthanasia. Blood was drawn through a cardiac puncture and collected in EDTA-containing vials. Mice were then perfused with 10 ml ice-cold PBS containing 2 mM EDTA; aortae were dissected and prepared for flow cytometry analyses; hearts were fixed, dehydrated, and embedded in paraffin or O.C.T. Compound (4583, Sakura Finetek USA) for histological analysis. All experiments were conducted according to a protocol approved by the Institutional Animal Care and Use Committee at the University of California, San Diego.

### Aorta single-cell suspension

The entire aorta from the aortic root to the iliac bifurcation (including ascending, arch, thoracic, and abdominal aorta) were used for flow cytometry analysis. Aortae were perfused with 10 mL cold PBS containing 2 mM EDTA. After removal of the surrounding fat and connective tissue, aortae were cut into several segments and digested in the aorta digestion enzyme solution^[1]^ [HBSS containing 250U /mL collagenase type XI (C7657, Sigma Aldrich), 120 U/mL hyaluronidase type I-s (H3506, Sigma Aldrich), 120 U/mL DNase I (DN25, Sigma Aldrich), and 450 U/mL collagenase type I (SCR103, Sigma Aldrich)] for 60 minutes at 37°C. The digested aortae were then passed through a 70 μM cell strainer (25-376, Olympus Plastics), and cells were pelleted by centrifugation and resuspended in RPMI containing 0.5% Poloxamer 188 (24040032, Thermo Fisher Scientific).

### Flow cytometry-based detection of inflammarafts

Aortic single-cell suspensions were incubated at 37°C for 30 minutes for cell surface recovery following enzymatic digestion and stained with Ghost Dye Red 780 Fixable Viability Dye (18452, Cell Signaling Technology). After neutral lipid staining with 1 μg/mL BODIPY 505/515 (D3921, Thermo Fisher Scientific), the single-cell suspensions were fixed with 4% PFA at 4°C for 20 minutes. Cells were blocked with an anti-CD16/CD32 antibody (553142, BD Biosciences) in PBS containing 2% BSA on ice for 30 minutes and then incubated with a mixture of BV650-conjugated anti-CD45 antibody (103151, Biolegend), PerCP/Cy5.5-conjugated anti-F4/80 antibody (123128, Biolegend), AF594-Conjugated anti-Cholera Toxin Subunit B antibody (C22842, Invitrogen), APC-conjugated anti-TLR4 (145406, BioLegend; detecting total TLR4 expression) antibody, and PE-conjugated anti-TLR4/MD-2 (Invitrogen, 12-9924-81; detecting TLR4 monomers) antibody for 60 minutes on ice. Cells were analyzed on a CytoFLEX Flow Cytometer (Beckman Coulter). Splenic F4/80-positive macrophages obtained from *Ldlr^−/−^*mice fed a standard laboratory diet were treated for 15 minutes with 100 ng/ml LPS (to induce transient TLR4 dimerization) and served as a positive control for the TLR4 dimerization assay. Non-stimulated splenic macrophages served as a negative control to standardize data obtained over time. The expression of TLR4 dimers was calculated from geometric mean fluorescence intensities of PE-conjugated TLR4/MD2 antibody (monomers) and APC-conjugated TLR4 antibody (total) as the percentage of dimers in total TLR4. TLR4 dimers and lipid rafts were then normalized to those in LPS-treated splenic F4/80-positive macrophages in each experiment to account for the inter-day assay variation.

### Immunofluorescence staining of the aortic root

Briefly, consecutive 10 μm-thick aortic cross-sections were collected with a Dakewe CT520 Research Cryostat. Sections were blocked with 3% FBS in PBS for 1 hour at room temperature and incubated with rat anti-F4/80 antibody (MF48000, Invitrogen) at 4 ℃ overnight. The sections were then incubated with anti-rat secondary antibody (A78947, Invitrogen) at room temperature for 2 hours, and counterstained with LipidTOX neutral lipid stain (H34476, ThermoFisher) for 30 min. Sections were mounted with DAPI-containing mounting media (P36931, ThermoFisher). Images were captured using a Leica Sp8 confocal microscope.

### LDL Oxidation

OxLDL was produced in vitro as previously reported.^[3]^ Briefly, human native LDL (36010, Lee BioSolutions) was extensively dialyzed against PBS to remove EDTA, and 0.1 mg/mL of LDL was incubated with 10 μM CuSO_4_ for 18 hours at 37 °C. Thiobarbituric acid reactive substances (typically, >30 nmol/mg in OxLDL) were measured to confirm LDL oxidation. OxLDL was concentrated to 1 mg/mL using a 100 kDa cutoff centrifugal concentrator (UFC810024, Millipore) and sterile filtered (0.22 μm). Endotoxin contamination was tested using a Pierce Chromogenic Endotoxin Quant Kit (A39553, Thermo Fisher), and only LDL preparations with endotoxin levels below 0.025 EU/mg protein were used in this study.

### Bone marrow-derived macrophages

BMDMs were cultured as previously described^[3]^. Briefly, bone marrow cells were isolated from femurs and tibias of 8-week-old *Apoa1bp^−/−^Ldlr^−/−^* mice^[12]^ and *Ldlr^−/−^* male mice and cultivated in L929-conditioned medium for one week. Cells were then replated in 6-well plates at a density of 1 million cells per well for a 24-hour resting period. Cells were incubated with 20 μg/mL OxLDL in DMEM supplemented with 5% lipoprotein-deficient serum. After the 24-hour incubation, cells were harvested for RNA isolation using the NucleoSpin RNA purification kit (740955.250, MACHEREY-NAGEL).

### RNA-seq

RNA samples were quantified and qualified using a high-sensitivity RNA screen tape kit (5067-5579, Agilent Technologies). Libraries of RNA samples were prepared using a Stranded mRNA Prep kit (20040534, Illumina) and sequenced on a NovaSeq XPlus sequencer (Illumina). Sequencing reads were quality-checked by fastqc^[23]^, cleaned by trim-galore^[**24**]^, mapped to the GRCm39 mouse genome using STAR^[25]^, and counted using featureCounts^[26]^. Differential gene expression analysis was performed using the DESeq2 R package^[27]^. Raw read counts were normalized using the DESeq2, and differential expression between conditions was estimated with the Wald test, incorporating Benjamini–Hochberg correction to control the false discovery rate (FDR). Genes were considered significantly differentially expressed if they exhibited an adjusted p-value < 0.05 and an absolute log2 fold-change greater than log2(1.5). To prioritize genes with both strong statistical support and substantial effect size, significant genes were independently ranked by adjusted p-value (ascending) and by absolute log2 fold-change (descending). Genes were subsequently ordered according to a composite score of the sum of these two rank values.

### Quantification of atherosclerotic lesions

Hearts were fixed and embedded in paraffin for the assessment of atherosclerosis as described previously.^[3]^ Briefly, consecutive 10 μm-thick aortic cross-sections were collected in 100 μm increments starting from the first appearance of the first leaflet of the aortic valve until the last leaflet. Sections were stained with modified Van Gieson staining to enhance the contrast between the intima and surrounding tissues. Images were acquired using a Nano Zoomer Slide Scanner. At each 100 μm increment, the lesion size and necrotic core size in each mouse were analyzed by computer-assisted morphometry (Image-Pro Plus 6.3, Media Cybernetics) by two investigators blinded to the study protocol. Lesion volumes were analyzed by calculating the area under the curve in each mouse. Average necrotic core percentages were analyzed by calculating the area under the curve / section numbers in each mouse.

### Measurements of cholesterol and triglycerides

Mice were fasted for 8 hours before euthanasia. Total cholesterol and triglyceride levels in plasma samples were quantified using a cholesterol/cholesteryl ester assay kit (ab65359, Abcam) and a triglyceride assay kit (ab65336, Abcam), respectively, according to manufacturer’s protocol.

### Statistical Analyses

Data are presented as mean ± SEM. N represents biological replicates. Data were analyzed for normality and equal variance using the Kolmogorov-Smirnov test. A two-sided Student’s *t*-test was employed to compare two groups. For multiple-group comparisons with one or two independent factors, one-way or two-way ANOVA was conducted with Tukey’s or Dunnett’s multiple comparison tests for normally distributed variables. All statistical analyses were performed using GraphPad Prism version 9.4.1. A p-value of less than 0.05 was considered statistically significant.

## Acknowledgments

The authors thank Dr Nicholas Webster for generously sharing access to a flow cytometer in his lab.

## Data availability statement

The raw sequencing data have been deposited in the NCBI SRA under accession PRJNA1377207. Other datasets generated during and/or analyzed during the current study are available from the corresponding author on reasonable request.

## Supplemental Materials

**Supplemental Figure S1.**
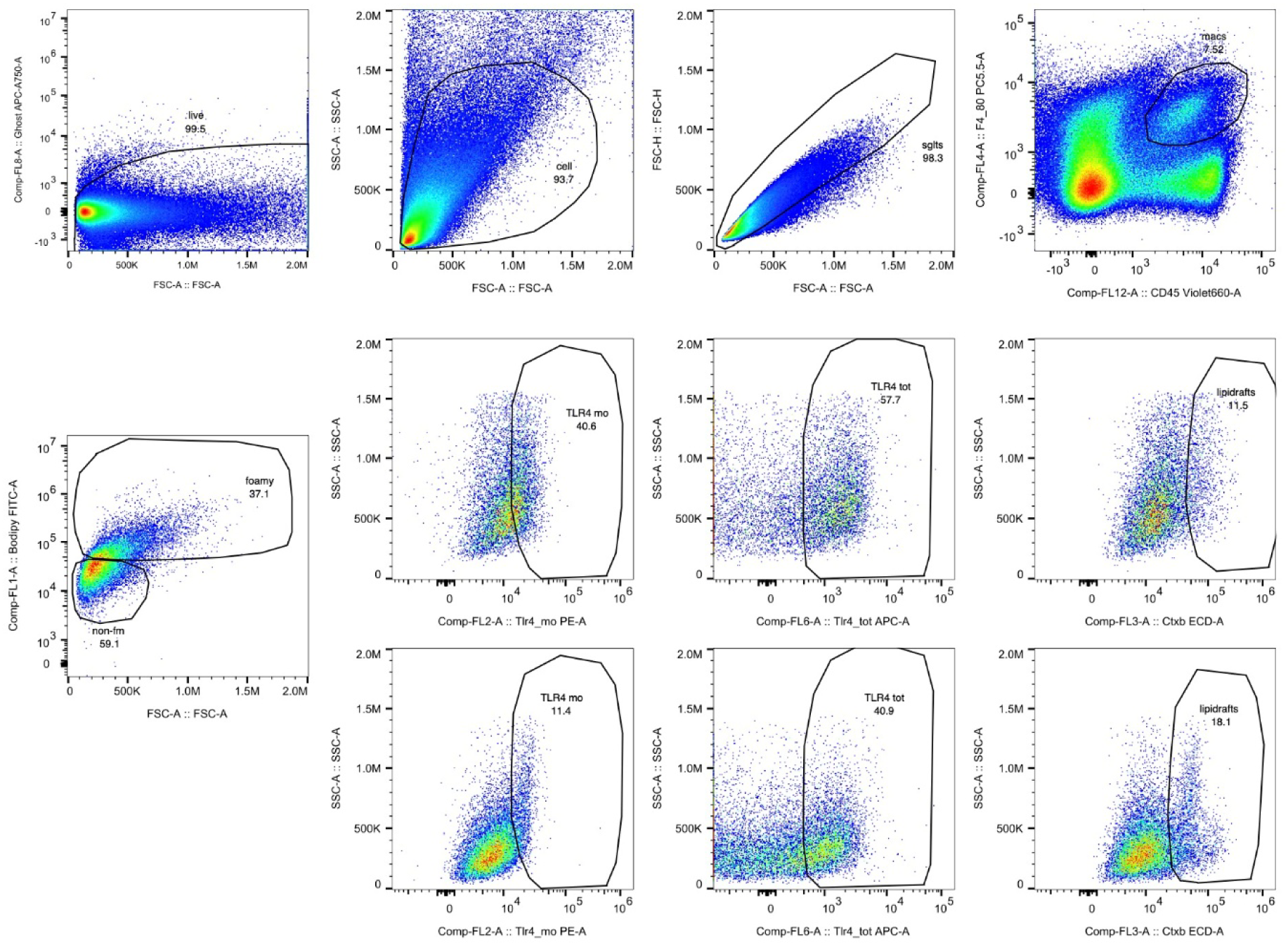
Gating strategy for flow cytometry analysis of expression of TLR4 dimers and lipid rafts. Aortic single-cell suspensions from *Apoa1bp^−/−^Ldlr^−/−^* and *Ldlr^−/−^* mice fed a 16-week high-fat diet were gated for BODIPY-high foamy and BODIPY-low non-foamy, CD45^+^ F4/80^+^ macrophages. The percentage of TLR4 dimers was calculated from geometric mean fluorescence intensities of PE-conjugated TLR4/MD2 antibody (monomers) and APC-conjugated TLR4 antibody (total). Lipid rafts were calculated from the geometric mean fluorescence intensity of AF594-conjugated cholera toxin B subunit.

**Supplemental Figure S2.**
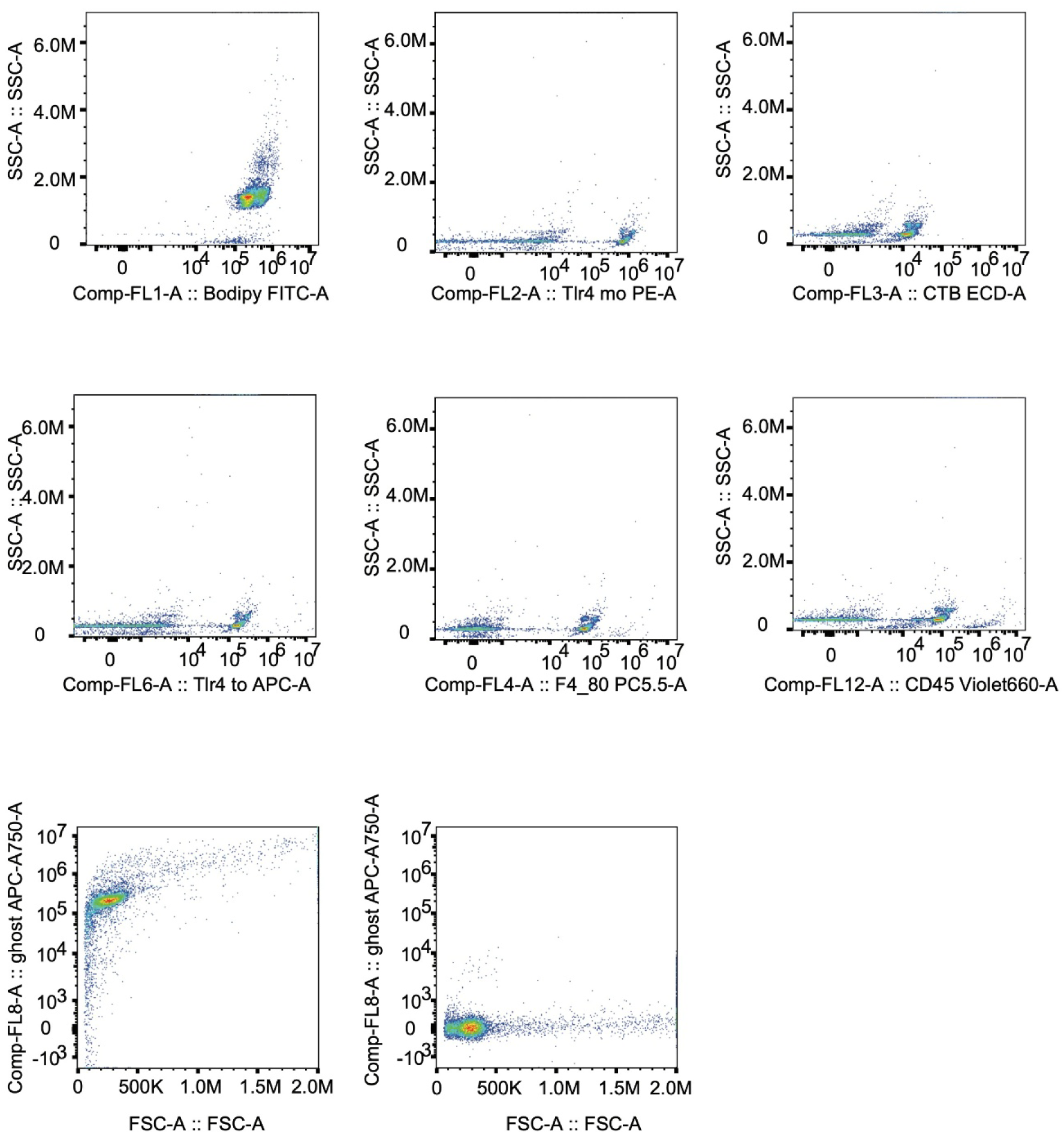
Single staining controls for compensation in flow cytometry.

## References

1 Kim, K. et al. Transcriptome Analysis Reveals Nonfoamy Rather Than Foamy Plaque Macrophages Are Proinflammatory in Atherosclerotic Murine Models. Circulation Research 123, 1127–1142 (2018). 10.1161/CIRCRESAHA.118.312804

2 Li, C. et al. AtheroSpectrum Reveals Novel Macrophage Foam Cell Gene Signatures Associated With Atherosclerotic Cardiovascular Disease Risk. Circulation 145, 206–218 (2022). 10.1161/CIRCULATIONAHA.121.054285

3 Navia-Pelaez, J. M. et al. Differential Expression of Inflammarafts in Macrophage Foam Cells and in Nonfoamy Macrophages in Atherosclerotic Lesions—Brief Report. Arteriosclerosis, Thrombosis, and Vascular Biology 43, 323–329 (2023). 10.1161/ATVBAHA.122.318006

4 Miller, Y. I., Navia-Pelaez, J. M., Corr, M. & Yaksh, T. L. Lipid rafts in glial cells: role in neuroinflammation and pain processing. J Lipid Res 61, 655–666 (2020). 10.1194/jlr.TR119000468

5 Dib, L. et al. Lipid-associated macrophages transition to an inflammatory state in human atherosclerosis, increasing the risk of cerebrovascular complications. Nature Cardiovascular Research 2, 656–672 (2023). 10.1038/s44161-023-00295-x

6 Anttila, V. et al. Genome-wide meta-analysis identifies new susceptibility loci for migraine. Nature Genetics 45, 912–917 (2013). 10.1038/ng.2676

7 Zhang, M. et al. Apolipoprotein A-1 binding protein promotes macrophage cholesterol efflux by facilitating apolipoprotein A-1 binding to ABCA1 and preventing ABCA1 degradation. Atherosclerosis 248, 149–159 (2016). 10.1016/j.atherosclerosis.2016.03.008

8 Matsuo, M. ABCA1 and ABCG1 as potential therapeutic targets for the prevention of atherosclerosis. J Pharmacol Sci 148, 197–203 (2022). 10.1016/j.jphs.2021.11.005

9 Ritter, M. et al. Cloning and characterization of a novel apolipoprotein A-I binding protein, AI-BP, secreted by cells of the kidney proximal tubules in response to HDL or ApoA-I. Genomics 79, 693–702 (2002). 10.1006/geno.2002.6761

10 Choi, S.-H. et al. Intracellular AIBP (Apolipoprotein A-I Binding Protein) Regulates Oxidized LDL (Low-Density Lipoprotein)-Induced Mitophagy in Macrophages. Arteriosclerosis, Thrombosis, and Vascular Biology 41, e82–e96 (2021). 10.1161/ATVBAHA.120.315485

11 Duan, M. et al. Mitochondrial apolipoprotein A-I binding protein alleviates atherosclerosis by regulating mitophagy and macrophage polarization. Cell Communication and Signaling 20, 60 (2022). 10.1186/s12964-022-00858-8

12 Schneider, D. A. et al. AIBP protects against metabolic abnormalities and atherosclerosis. J Lipid Res 59, 854–863 (2018). 10.1194/jlr.M083618

13 Kim, Y. S. et al. AIBP controls TLR4 inflammarafts and mitochondrial dysfunction in a mouse model of Alzheimer’s disease. Journal of Neuroinflammation 21, 245 (2024). 10.1186/s12974-024-03214-4

14 Woller, S. A. et al. Inhibition of Neuroinflammation by AIBP: Spinal Effects upon Facilitated Pain States. Cell Reports 23, 2667–2677 (2018). 10.1016/j.celrep.2018.04.110

15 Hu, Y. H. et al. WDFY1 mediates TLR3/4 signaling by recruiting TRIF. The EMBO Reports 16, 447–455 (2015). 10.15252/embr.201439637

16 Dabravolski, S. A. et al. Purinergic receptors in atherosclerosis: implications for pathophysiology and therapeutic strategies. Journal of Physiology and Biochemistry (2025). 10.1007/s13105-025-01108-4

17 Di Nubila, A., Dilella, G., Simone, R. & Barbieri, S. S. Vascular Extracellular Matrix in Atherosclerosis. International Journal of Molecular Sciences 25, 12017 (2024).

18 De Meyer, G. R. Y., Zurek, M., Puylaert, P. & Martinet, W. Programmed death of macrophages in atherosclerosis: mechanisms and therapeutic targets. Nature Reviews Cardiology 21, 312–325 (2024). 10.1038/s41569-023-00957-0

19 Puylaert, P., Zurek, M., Rayner, K. J., De Meyer, G. R. Y. & Martinet, W. Regulated Necrosis in Atherosclerosis. *Arteriosclerosis*, Thrombosis, and Vascular Biology 42, 1283–1306 (2022). 10.1161/ATVBAHA.122.318177

20 Spann, N. J. et al. Regulated accumulation of desmosterol integrates macrophage lipid metabolism and inflammatory responses. Cell 151, 138–152 (2012). 10.1016/j.cell.2012.06.054

21 Yerly, A. et al. Sex-specific and hormone-related differences in vascular remodelling in atherosclerosis. European Journal of Clinical Investigation 53, e13885 (2023). 10.1111/eci.13885

22 Man, J. J., Beckman, J. A. & Jaffe, I. Z. Sex as a Biological Variable in Atherosclerosis. Circulation Research 126, 1297–1319 (2020). 10.1161/CIRCRESAHA.120.315930

23. 23 Andrews, S. FastQC: A Quality Control Tool for High Throughput Sequence Data. Available online at: http://www.bioinformatics.babraham.ac.uk/projects/fastqc/ (2010).

24 Felix Krueger, F. J., Phil Ewels, Ebrahim Afyounian, Michael Weinstein, Benjamin Schuster-Boeckler, Gert Hulselmans, & Sclamons. FelixKrueger/TrimGalore: v0.6.10 - add default decompression path. Zenodo (2023). 10.5281/zenodo.7598955

25 Dobin, A. et al. STAR: ultrafast universal RNA-seq aligner. Bioinformatics 29, 15–21 (2012). 10.1093/bioinformatics/bts635

26 Liao, Y., Smyth, G. K. & Shi, W. featureCounts: an efficient general purpose program for assigning sequence reads to genomic features. Bioinformatics 30, 923–930 (2013). 10.1093/bioinformatics/btt656

27 Love, M. I., Huber, W. & Anders, S. Moderated estimation of fold change and dispersion for RNA-seq data with DESeq2. Genome Biology 15, 550 (2014). 10.1186/s13059-014-0550-8

